# An intranasally administrated SARS-CoV-2 beta variant subunit booster vaccine prevents beta variant viral replication in rhesus macaques

**DOI:** 10.1101/2021.10.19.464990

**Authors:** Yongjun Sui, Jianping Li, Hanne Andersen, Roushu Zhang, Sunaina Kiran Prabhu, Tanya Hoang, David Venzon, Anthony Cook, Renita Brown, Elyse Teow, Jason Velasco, Laurent Pessaint, Ian N. Moore, Laurel Lagenaur, Jim Talton, Matthew W. Breed, Josh Kramer, Kevin W. Bock, Mahnaz Minai, Bianca M. Nagata, Hyoyoung Choo-Wosoba, Mark G. Lewis, Lai-Xi Wang, Jay A. Berzofsky

**Author notes:** Correspondence to: Yongjun Sui, Vaccine Branch, National Cancer Institute, National Institutes of Health, 41 Medlars Drive, Bethesda, MD 20892 USA; Jay A. Berzofsky, Vaccine Branch, National Cancer Institute, National Institutes of Health, 41 Medlars Drive, Bethesda, MD 20892 USA. **Competing interests** Authors declare no competing interests.

## Abstract

Emerging of SARS-CoV-2 variants and waning of vaccine/infection-induced immunity poses threats to curbing the COVID-19 pandemic. An effective, safe, and convenient booster vaccine will be needed. We hypothesized that a variant-modified mucosal booster vaccine might induce local immunity to prevent SARS-CoV-2 infection at the port of entry. The beta-variant is hardest to cross-neutralize. Herein we assessed the protective efficacy of an intranasal booster composed of beta variant-spike protein S1 with IL-15 and TLR agonists in previously immunized macaques. The macaques were first vaccinated with Wuhan strain S1 with the same adjuvant. One year later, negligibly detectable SARS-CoV-2-specific antibody remained. Nevertheless, the booster induced vigorous humoral immunity including serum- and bronchoalveolar lavage (BAL)-IgG, secretory nasal- and BAL-IgA, and neutralizing antibody against the original strain and/or beta variant. Beta-variant S1-specifc CD4^+^ and CD8^+^ T cell responses were also elicited in PBMC and BAL. Following SARS-CoV-2 beta variant challenge, the vaccinated group demonstrated significant protection against viral replication in the upper and lower respiratory tracts, with almost full protection in the nasal cavity. The fact that one intranasal beta-variant booster administrated one year after the first vaccination provoked protective immunity against beta variant infections may inform future SARS-CoV-2 booster design and administration timing.

## INTRODUCTION

Emergence of novel SARS-CoV-2 variants of concern (VOC) threatens the efforts to curb the COVID-19 pandemic. Some variants demonstrated significantly reduced neutralization sensitivity to sera from convalescent and vaccinated individuals. A recent study assessed the cross-reactive neutralizing responses to different variants including B.1.1.7 (Alpha), B.1.351 (Beta), P.1 (Gamma), B.1.429 (Epsilon), B.1.526 (Iota), and B.1.617.2 (Delta) in mRNA-1273 vaccinated individuals, and found that the beta variant had the lowest antibody recognition ^1^. To date, the beta variant seems to be one of the most resistant variants to convalescent and vaccinated sera ^1,2^. This variant was first detected in South Africa in October 2020 from samples collected at Eastern Cape Province in early August ^3,4^. Since then, it quickly spread within South Africa and to the other parts of the world. By December 2020, it spread all over the world, accounted for 87% of viruses sequenced in South Africa, and became the dominant strain in Zambia ^4,5^. Multiple mutations were found in this variant with K417N, E484K, N501Y as key substitutions ^6^. It had 5-fold enhanced affinity to ACE2 compared to the original virus ^7^, and a 3.5-fold ^8^, and a 10-fold ^2^ reduction in neutralization ability in convalescent and vaccinated individuals. Two studies have showed that the beta variant can partially or completely escaped three classes of therapeutically relevant antibodies and the convalescent sera ^9,10^.

Meanwhile, waning immunity after vaccination has led to a gradual decline of vaccine efficacy against SARS-CoV-2 infections ^11-13 14,15^. Recently, more SARS-CoV-2 breakthrough infections in vaccinated individuals, and resurgence of SARS-CoV-2 cases in some countries have been observed. Based on the previous experience with other coronaviruses and the current situation, an extra booster with the original Pfizer-BioNTech mRNA vaccine after 6-months of the first vaccination has been authorized in some countries among individuals with older age, high risk for severe COVID-19, or high risk for SARS-CoV-2 infections due to occupational or institutional exposure ^16^. For the general population, it is anticipated that a booster, ideally targeting circulating viral variants, will be needed, when the immunity induced by the original vaccine cannot provide adequate protection against the circulating viral variants ^17^. Since a large number of individuals have been vaccinated with the vaccines comprised of antigens from the SARS-CoV-2 original Wuhan strain, data on immunogenicity and protective efficacy of a variant booster to vaccinees, who have previously received the original vaccines, would be urgently needed ^18,19^. Recent studies have shown that intranasal administration of different platforms of SARS-CoV-2 vaccines induce protective immunity in preclinical animal models ^20-24^.

Herein we performed a proof-of-concept study to test the immunogenicity and efficacy of an adjuvanted SARS-CoV-2 beta variant subunit booster in rhesus macaques that were vaccinated with the same vaccine platform except that the spike protein S1 was from the original Wuhan strain. We found that one year after the first vaccination, almost no detectable immunity was present in these macaques. However, an intranasal booster with the adjuvanted beta variant S1 subunit vaccine induced vigorous humoral and cellular immunity against both original and beta variant antigens. Most importantly, secretory IgA responses against S1 from both the original Wuhan strain and the beta variant were detected in the nasal cavity, which was consistent with the almost full protection we observed against the beta variant in the nasal cavity after viral challenge. Whether this mucosal vaccine can protect against viral transmission, and whether the mucosal IgA response is responsible for the protection in the nasal cavity merits further investigation. Iimportantly, our data showed that the one-year intranasal booster with beta variant S1 protein reinvigorated SARS-CoV-2-specific immune responses and led to significant protection against beta variant challenge. This study may provide important information regarding the timing of booster immunizations and the type of antigens included in the booster.

## RESULTS

### Robust systemic and mucosal humoral responses against S1 from the original Wuhan strain and beta variant were elicited after intranasal variant booster

In this study, we took advantage of five Indian rhesus macaques that had been vaccinated one year earlier with S1 protein from the original Wuhan strain (Supplementary Table 1). The vaccine included 100 μg of S1 and CP15 adjuvant, which was composed of IL-15 and TLR agonists (CpG and Poly I:C) incorporated in PLGA nanoparticles as used in our previous study ^20^. The macaques were first primed with the vaccine intramuscularly (IM) at week 0, and then boosted with the same vaccine intranasally (IN) at week 3 and week 6 (Figure 1). 100 μg of S1 per dose was used based on our previous HIV and SARS-CoV-2 vaccine studies ^20,25^. S1 with sequence of the original Wuhan strain was used in the first three vaccinations. When evaluating the S1-specific IgG antibody responses, we found that this vaccine regimen induced a moderate level of humoral immune responses in serum and BAL fluid (Figure 2a). Compared to the ID50 of 25, 209 induced by IM-primed and -boosted alum-adjuvanted subunit vaccine ^20^, the peak median serum ID50 was only 945 (Figure 2a). Moreover, the vaccine-induced immunity also waned with time. After one year, the IgG responses in the vaccinated animals were comparable to those of the naïve controls (Figure 2a).

**Figure 1.**
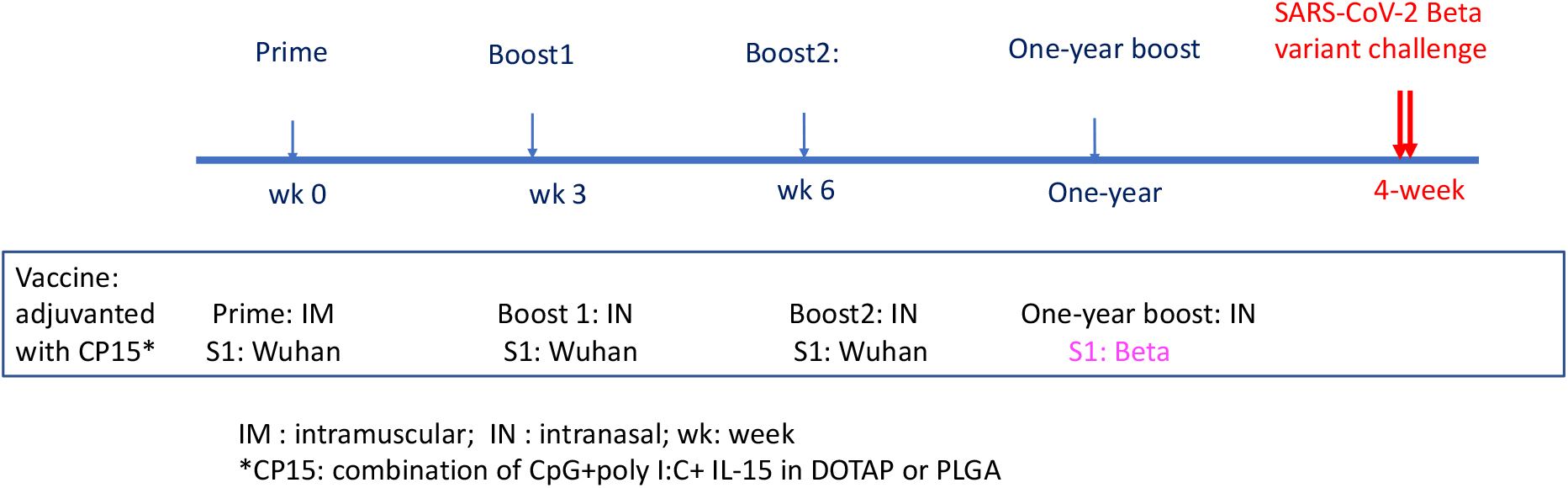
Schematic diagram of vaccination and viral challenge.

**Figure 2.**
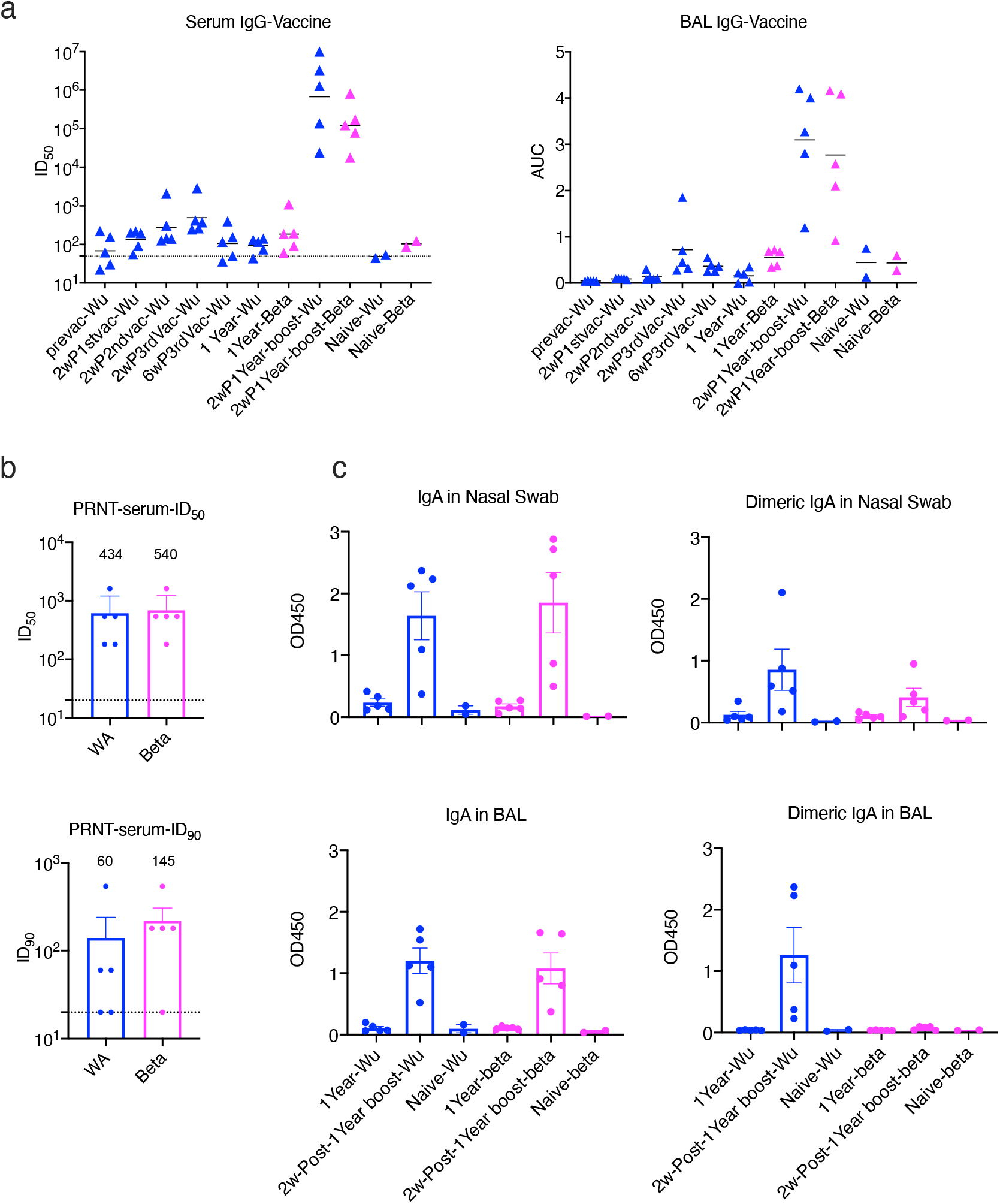
Humoral immune responses against SARS-CoV-2 spike protein 1 (S1) in vaccinated macaques. (a). the kinetics of S1-specific binding IgG titers in serum and BAL. Bars indicate geometric means of ID50 and means of AUC. (b). PRNT titers in the serum samples of the vaccinated animals at 2-week after one-year boost. Geometric mean + geometric SD are shown. (c) S1-specific IgA and dimeric IgA responses in nasal swabs (NS) and BAL samples. WA: WA1/2020 D614G SARS-CoV-2 strain; Wu: Wuhan original strain; Beta: B.1.351 variant. The dashed lines indicate the detection limits. Data are shown as mean + SEM. Blue color indicates the S1 protein or the virus from Wuhan or WA strain, and magenta color indicates from beta variant.

We then gave the animals one intranasal booster with S1 from beta variant adjuvanted with CP15 in DOTAP nanoparticles. After the booster, significant anamnestic responses were elicited. The Log ID50 of serum IgG titer reached 5.83 for the original Wuhan strain, and 5.08 for the beta variant, compared to the highest IgG titer of 2.77 logs at 2 weeks post the 3^rd^ vaccination one year earlier (Figure 2a). The booster also led to the induction of a substantial increase of mucosal IgG against both the original Wuhan strain and the beta variant S1 in BAL (Figure 2a).

High titers of live virus neutralization antibody (Nab) responses against both WA1/2020 D614G SARS-CoV-2 (WA) strain and the beta variant were detected in the serum. The geometric mean titers (GMT) of Nab were 434 and 540 for ID50, and 60 and 145 for ID90 for the WA strain and the beta variant, respectively (Figure 2b). Given the fact that the beta variant has been the most difficult strain to neutralize so far ^1^, boosting with beta variant S1 might account for this improvement and suggest the potential benefit of switching antigens from the original WA strain to a variant. It is noteworthy that boosting with the variant S1 still induced a strong anamnestic response against the original priming Wuhan S1.

IgA and dimeric IgA responses in bronchoalveolar lavage (BAL) and nasal swabs were also examined, as IgA, especially dimeric IgA, displays high binding affinity to pathogens, and thus is more potent at preventing mucosal pathogen infections ^26,27^. Right before the one-year booster, no S1 (original or beta variant)-specific IgA, or dimeric IgA responses were detected, and the antibody titers were comparable to the basal levels of naïve animals (Figure 2c). Consistent with IgG and neutralization responses, the one-year booster enhanced IgA responses in nasal swab and BAL samples with similar antibody titers against S1 from the original strain and the beta variant (Figure 2c). However, dimeric IgA responses against beta variant were not induced in BAL samples, whereas dimeric IgA responses were observed in BAL against the Wuhan strain and in nasal swabs against both strains (Figure 2c).

Overall, our results showed that the one-year booster induced robust S1-specific antibody responses in serum and BAL, including potent neutralizing antibody responses in peripheral blood. Most importantly, mucosal IgA responses were induced in nasal swabs and BAL that were comparable against both the original priming Wuhan strain and the beta variant, except dimeric IgA responses against beta variant in BAL.

### Correlations among different types of antibody responses

We next assessed the spearman correlations among different types of antibody responses. First, PRNT titers against the WA strain and beta variant did not correlate with each other (Supplementary Table 2), consistent with the fact that these two viruses have different neutralization profiling. Neutralization against one strain does not guarantee the neutralization of the other. Interestingly, the serum IgG responses against S1 from the original Wuhan strain did not correlate with any of the other antibody measurements, including serum IgG titer against the beta variant (Supplementary Table 2). In contrast, mucosal antibody responses, including S1-specific BAL IgG, IgA and dimeric IgA responses in nasal swabs, IgA responses in BAL, showed correlations or trends of correlations between the original Wuhan strain and beta variant (Supplementary Table 2 and Supplementary Figure 2). Moreover, the serum IgG titers against the beta variant were positively correlated (or showed trends of correlations) with BAL IgG and IgA, serum PRNT, and nasal dimeric IgA responses (Supplementary Table 2 and Supplementary Figure 2). These correlations suggested that the repertoire of systemic and mucosal humoral response against the original Wuhan strain and the beta variant were different after administration of the beta variant S1 one-year booster, even though the geometric mean titers against the Wuhan strain and beta variant were generally similar.

### Variant S1-specific cellular responses were induced after the one-year booster

The vaccine-induced S1-specific T cell responses in PBMC and BAL samples of the vaccinated animals were evaluated by intracellular cytokine staining. S1-specific type 1 helper T cell responses (Th1) and CD8^+^ T cell that secrete tumor necrosis factor (TNF)-α, and/or interferon (IFN)-γ were induced after the first vaccination (Figure 3). Though the responses were persistent in most of the vaccinated animals, no further enhancement of the responses was observed after the second and third vaccinations. For CD8^+^ T cell responses, especially the responses in PBMC, we observed a declining trend with each vaccination (less so in BAL). This raises the concern that extensive boosters in a short period of time might burn out the SARS-CoV-2 -specific T cell responses. Nevertheless, the responses waned to under the detection limit in most of the animals after one year. After the administration of the one-year beta-variant booster, the S1-specific CD8^+^ T cell responses were successfully recalled in all 5 PBMC samples and CD4^+^ responses in 4/5 (Figure 3). Even though the route of the one-year booster was intranasal, S1-specific CD4^+^ T cells were induced only in 3 BAL samples, and CD8^+^ T cells in only two. One possibility could be the migration of antigen-specific T cell to the nasal cavity, while the other could be antigen-specific T cell exhaustion caused by multiple doses of vaccine. Both possibilities warrant further investigation.

**Figure 3.**
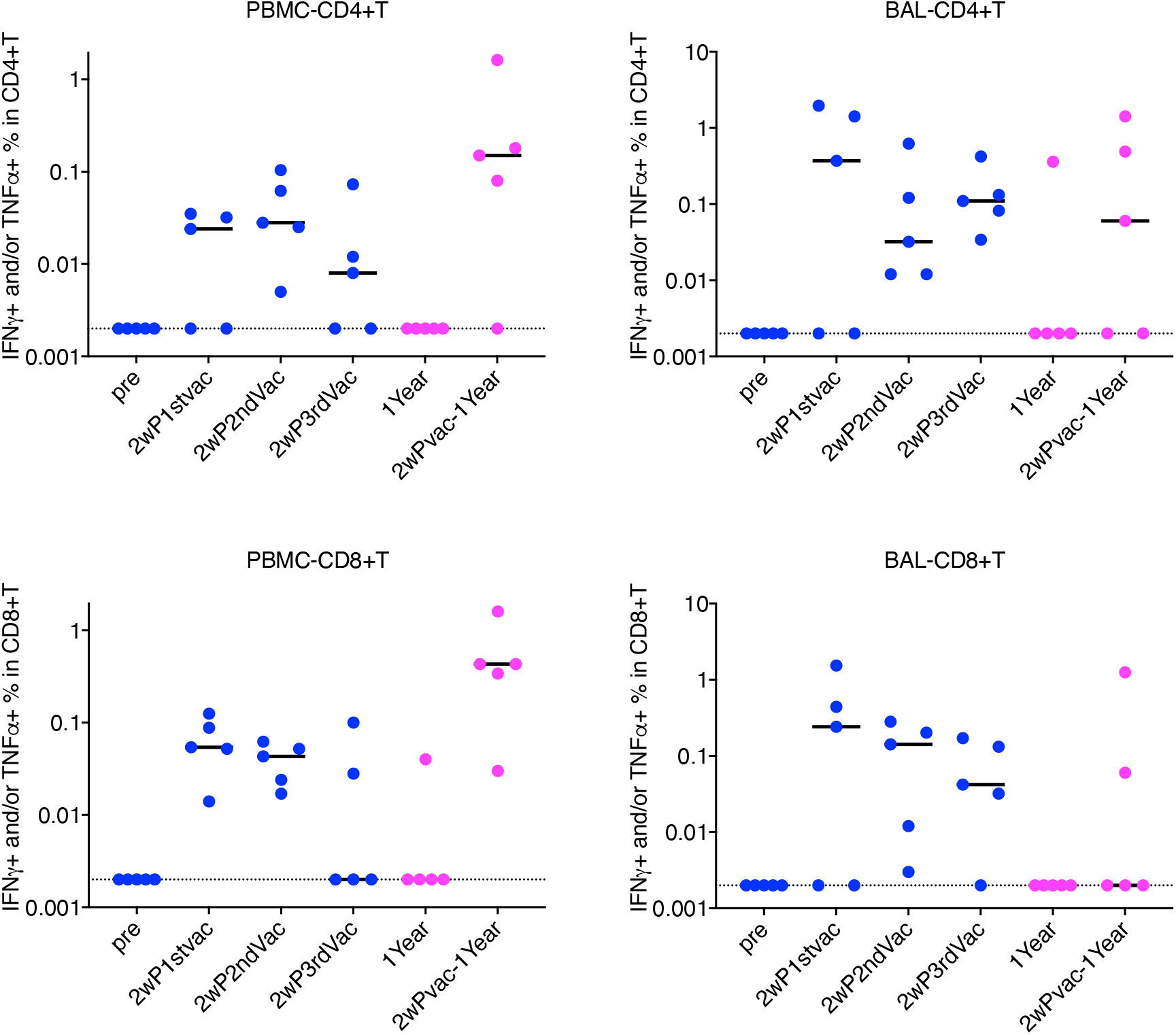
T cell responses against SARS-COV-2 spike protein 1 (S1) in PBMC and BAL samples of the vaccinated macaques. (a-b) The frequencies of IFNγ and/or TFNα-producing CD4^+^ and CD8^+^ T cells were stained and measured after stimulation with S1 for 18 hrs in PBMC and BAL samples. Dashed lines indicate the detection limits. Bars indicate medians. Blue color indicates the S1 protein or the virus from Wuhan or WA strain, and magenta color indicates from beta variant.

As the frequencies of antigen-specific T cell responses were low, we further assessed the kinetics of total Th1 and Th2 subsets after stimulation with Phorbol 12-myristic 13-acetate (PMA) and ionomycin. There were no significant alterations after the first three vaccinations in the prior year (Supplementary Figure 1). However, the one-year boost resulted in sharp increase of Th1 responses in PBMC while the Th2 responses did not change (Supplementary Figure1).

### Vaccinated animals demonstrated significant protection in BAL, and almost full protection in nasal swabs against SARS-CoV-2 beta variant replication

To test the protective efficacy against SARS-CoV-2 beta variant, 5 vaccinated and 5 naïve macaques were challenged with 1.0×10^5 TCID50 SARS-CoV-2 beta variant (isolate beta variant B.1.351, in-house generated stock from BEI Resources, NR-54974) through intranasal (1mL) and intratracheal (1mL) routes 4 weeks after the last vaccination. Viral tissue culture infectious dose 50 titers (TCID50) were measured in the collected nasal swab and lung BAL samples. Replicating viruses were detected in both nasal swabs and BAL samples of all five naïve animals, indicating that the viral inoculation was successfully delivered and propagated in the upper and lower airways (Figure 4a). It is worth mentioning that the inoculation of SARS-CoV-2 beta variant led to prolonged detection of replicating virus in the nasal turbinate of the naïve animals. High levels of viral replication were present in all five naïve animals at day 7 post virus challenge. In contrast, the vaccinated animals demonstrated almost full protection in nasal swabs: only one animal showed a small blip at day 2 post viral challenge, while four other vaccinated animals were free of replicating virus during the 7-days post-challenge period (Figure 4a). The vaccinated group showed significant reduction of viral replication in both nasal turbinate and lungs compared to naïve controls, based on the area under the curves over all time points (Figure 4b). A future transmission study is needed to test whether this mucosal booster can prevent transmission.

**Figure 4.**
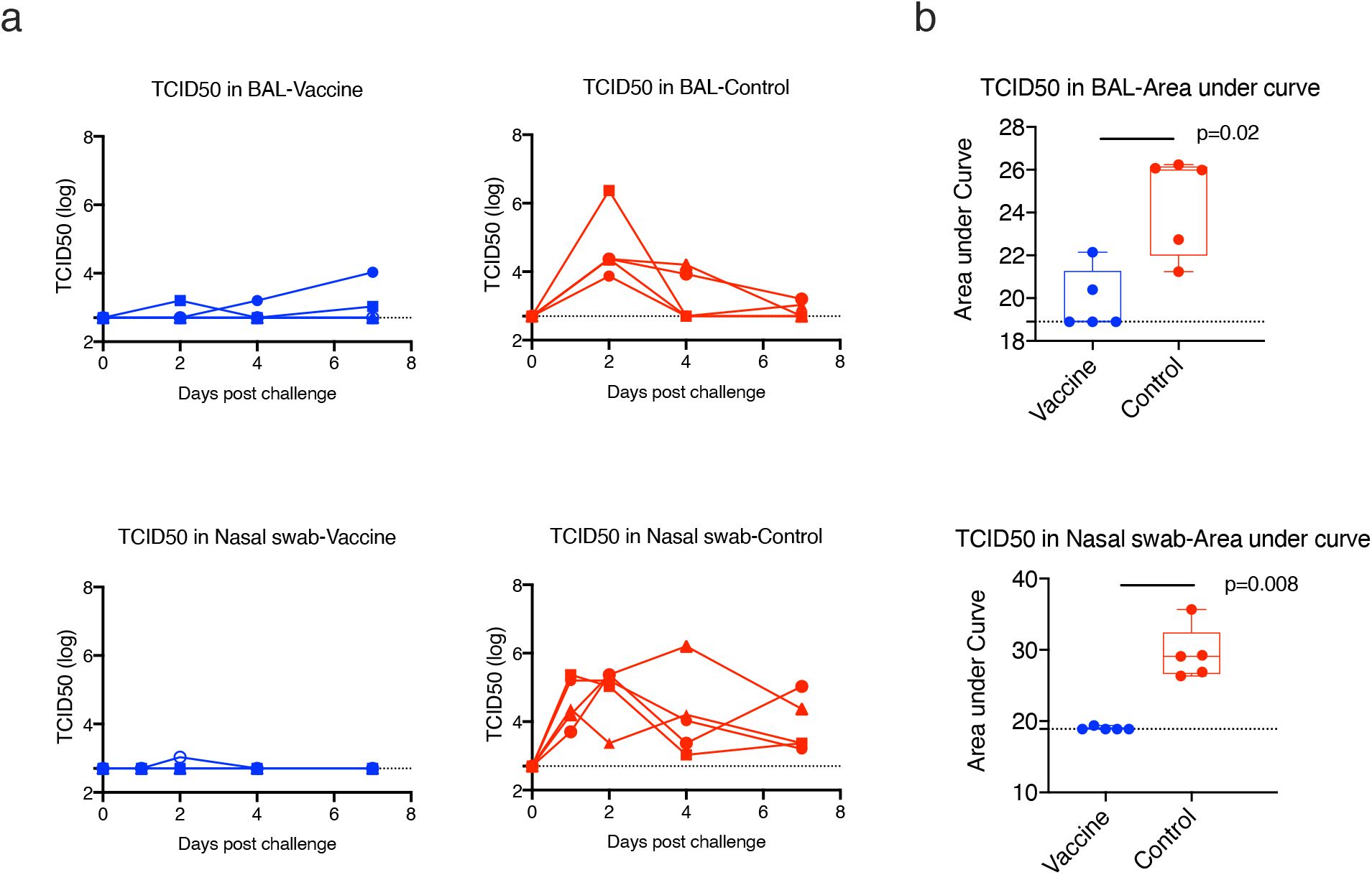
Viral burden in the nasal swabs (NS) and BAL samples after SARS-CoV-2 beta variant intranasal and intratracheal challenges. (a). TCID50 titer of the viral burdens in nasal swabs (NS) and BAL samples of individual animals (n=5 in the vaccine group and n=5 in the control group). (b). Area under curve (AUC) over time after challenge was calculated for each animal, representing total viral burdens. The total viral burdens were compared between vaccine and control groups in NS and BAL. Dashed lines indicate the detection limits. Box and whiskers with min to max were shown in the graph.

### Histopathology in the lungs after viral infection confirmed the protection in lungs

As reported in the previous study ^20^, the mucosal vaccine is safe. Throughout the whole course of this study, we did not observe any adverse effects in the vaccinated animals. When the animals were necropsied on day 7, sections of lung were evaluated immunohistochemically for SARS-CoV-2 virus antigen and histologically for the presence of SARS-CoV-2 -associated inflammation. None of the 5 vaccinated animals demonstrated immunoreactivity to viral antigens, while virus antigens were detected in the lung sections of the 4 out of 5 animals in the control group (Figure 5a-b). Predominantly perivascular to interstitial inflammation was observed in the control group. An inflammation score was given to each animal blindly by a certified pathologist based on the evaluation of lung infiltration collected at the time of necropsy at day 7 post SARS-CoV-2 challenges (Supplementary Table 3). The inflammation score was slightly more severe in the control group compared to the vaccinated group (Figure 5c), suggesting the vaccination prevented inflammation in the lungs.

**Figure 5.**
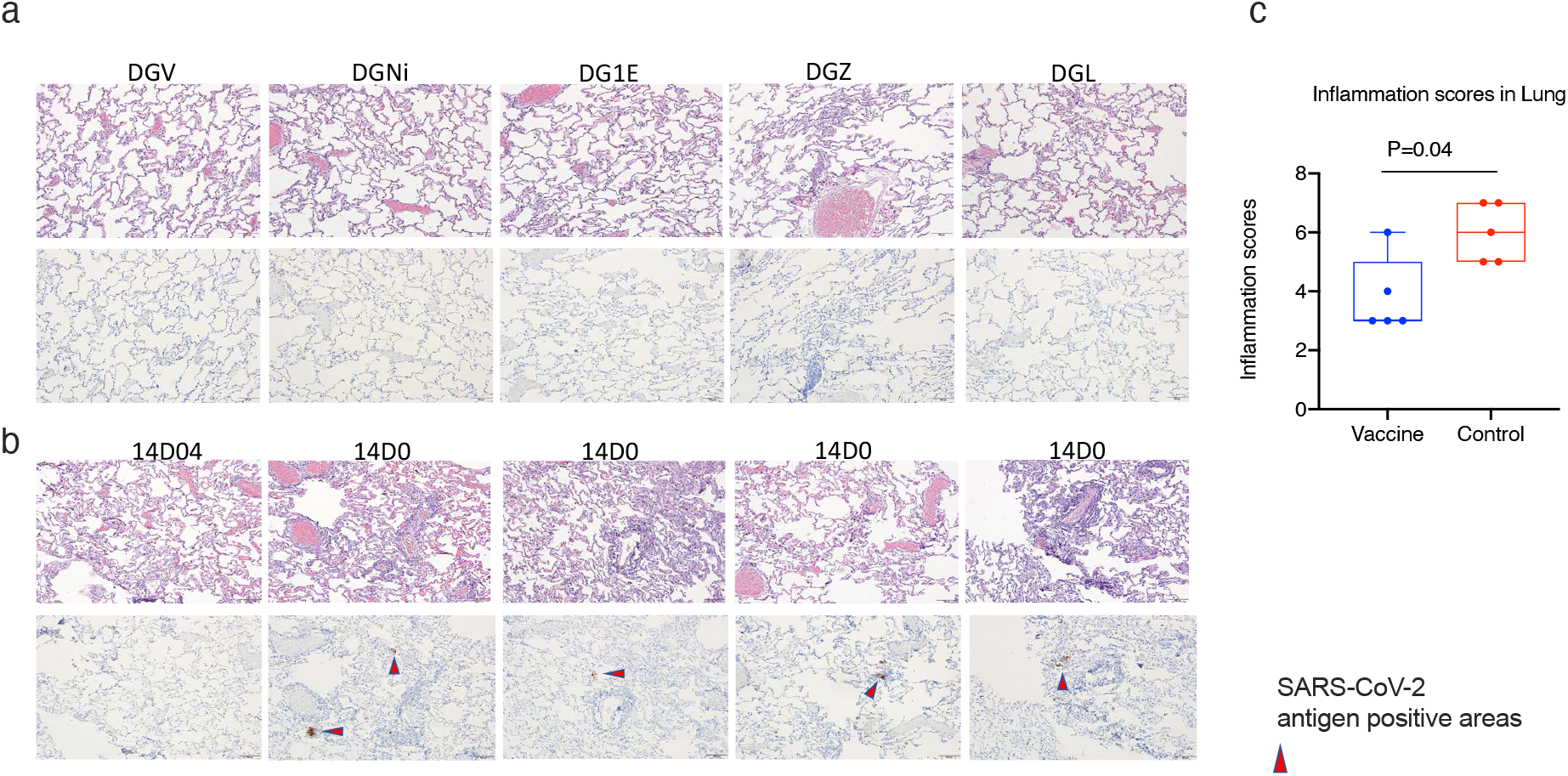
Histopathology in the lungs at day 7 post SARS-CoV-2 challenge. H&E and immunohistochemistry to detect SARS-COV-2 antigens were performed in the vaccinated(a) and naïve (b) animals. The upper rows of a-b were H&E staining, while the lower rows of a-b were immunohistochemistry of SARS-CoV-2 detection. All images 10x (scale bar= 100um). c). inflammation scores in the lung were compared between the vaccinated and naïve groups. Mann-Whitney test was used for comparison. Box and whiskers with min to max were shown in the graph.

## DISCUSSION

An additional booster vaccine is likely needed to curb the resurgence of SARS-CoV-2 cases. We demonstrated here that the one-year beta variant mucosal booster given intranasally elicited high quality immune responses and mediated protections against subsequent SARS-CoV-2 beta variant viral challenge in rhesus macaques. Notably, the protection in the upper respiratory tract was better than in the lower respiratory tract, which is different from most of the systemic vaccines ^28-31^. The nearly full protection against viral replication in the nasal cavity is especially encouraging, indicating its potential to prevent viral spread and transmission. The nasal mucosa is the first site of infection, so the local immunity might be able to abort viral replication here before it disseminates systemically and may also prevent spread to other individuals. Indeed, we found that high titers of mucosal IgA responses against both original and variant spike proteins were induced in the nasal mucosa, which might account for the efficient clearing of the virus in situ. These findings show the promise of a nasal mucosal vaccine as a booster rather than another systemic (IM) vaccine dose.

Waning immunity over time after vaccination/infection is contributing significantly to the resurgence of SARS-CoV-2 cases ^14,15,32^. Though the immune correlates of protection have not been fully established, neutralizing antibody (Nab) responses are believed to be one of the major protective mechanisms ^33-35^. To evaluate the durability, one study found that the half-life of Nab was biphasic, with a rapid initial decline over 61 days, and then a more gradual tapering after the first 2 months out to 104 days ^36^, while the other study found that Nab exhibited a bi-phasic decay with an extended half-life of >200 days ^37^. Though prolonged humoral and cellular immunity up to 10 months or one year has been reported in SARS-CoV-2-convalescent individuals ^38-40^, the durability of the protective immunity against SARS-CoV-2 infection remains unknown.

The emergence of SARS-CoV-2 variants of concern might partially account for the reported decreased vaccine effectiveness after 6 months ^12,41^. These variants either have high infectious potency or evade the immunity induced by SARS-CoV-2 infection or vaccination. The beta variant has the greatest immune evasive capacity among the widespread variants detected to date ^1^. In this study, we have switched the S1 from original Wuhan strain to that of the beta variant, which led to successful elicitation of systemic and mucosal immune responses against both the original strain and the beta variant, and most importantly mediated protection against subsequent SARS-CoV-2 beta variant challenge. Incorporating S1 from the beta variant into the booster vaccine might account for the observed robust protection.

A dramatic increase in antibody titers after the one-year booster was observed (more than 3 log of increase compared to the highest titers one year before for serum IgG titers). This is consistent with what we have found in a previous study, where the booster at 4 months induced much higher quality SARS-CoV-2 specific immune responses than the booster at 3 weeks did ^20^. It appears that the longer interval between the booster and the previous vaccinations enhances the immune responses. Similar phenomena were reported in AstraZeneca (AZ) and inactivated vaccine trials, as well as in the standard hepatitis B viral vaccine regimen. In the AZ trial, a longer prime-boost interval (>12 weeks) led to higher vaccine efficacy compared to shorter interval (<6 weeks) ^42^. In an inactivated vaccine trial, 6 or more months between the second and third vaccinations also induced a remarkable increase in antibody levels compared to a 4-week interval ^43^. Thus, these studies should be taken into consideration when deciding the timing of an additional booster.

The CP15 adjuvanted vaccine described here was not very effective as a prime vaccine. It did not induce robust immune responses compared to an alum adjuvanted vaccine ^20^. One year after the first vaccination, no virus-specific humoral or cellular immunity was detected. Nevertheless, the one-year booster elicited high quality immune responses, and mediated protection against subsequent beta variant challenge, which suggested that the vaccinations in the prior year generated persistent SARS-CoV-2 specific immune memory. Though the humoral and cellular immune responses waned to undetectable levels after one year, the immune memory persisted, which facilitated the later recall responses, when boosted. Moreover, our data suggest that a weaker variant-modified booster vaccine might be sufficient to induce protective immunity in previously vaccinated hosts. These findings may help guide future prime-boosting regimens for COVID-19.

## METHODS

### Animals

10 Indian-origin adult male rhesus macaques (*Macaca mulatta*), 3-8 years old, were enrolled in the study. The animals tested seronegative for cercopithecine herpesvirus 1, SIV, simian type-D retrovirus, simian T lymphotropic virus type 1, and SARS-CoV-2 prior to study assignment.

### Vaccine design and inoculation

Five macaques were included in the vaccine group, while five were in the SARS-CoV-2-naive control group. The five naïve control animals had been exposed to HIV envelope protein/glycopeptide vaccination more than one year before. The five macaques in the vaccine group were primed at Week 0 (administrated IM) and boosted at Week 3 (administered IN) and Week 6 (administered IN) with SARS-CoV-2 S1 protein (WA strain) with alum or CP15 adjuvant in PLGA nanoparticles. The CP15 adjuvant was composed of 200 μg per dose of D-type CpG oligodeoxynucleotide, 1 mg per dose of Poly I:C (InvivoGen), and 200 μg per dose of recombinant human IL-15 (Sino Biological). One year later, a boost was given to the remaining five animals with S1 protein from the beta variant adjuvanted with CP15. 100 μg of recombinant SARS-CoV-2 (2019-nCoV) spike S1 protein (Cat: 40591-V08H and 40591-V08H10, Sino Biological, endotoxin level: <0.001U/μg) was used per dose. S1 protein and CP15 were formulated in nanoparticles in PLGA (Alchem Laboratories) for the first 2 doses and the last (one-year) boost was in DOTAP (100 μl per dose; Roche). For immunization, the CP15 adjuvanted vaccine was given either intramuscularly in 1ml of volume, or intranasally in a volume of 50 µl per nostril, while the animals were anesthetized. After vaccination, blood, nasal swab and BAL fluid samples were collected at the times noted and analyzed.

### Nasal swab and BAL sample collection

Nasal secretions were collected and stored at -80°C after either using cotton-tipped swabs and then in 1 ml of PBS buffer containing 0.1% BSA, 0.01% thimerosal, and 750 Kallikrein inhibitor units of aprotinin ^25^ for pre-challenge stage, or using Copan flocked swabs and in virus transport medium for post-challenge stage. BAL samples were collected as described before ^20^. Briefly, while the animals were under anesthesia, up to 10 mL/kg of sterile saline were instilled into and sucked out of the lungs. Large pieces were removed by passing through a 100 μm cell strainer (pre-challenge). The BAL fluid was collected after centrifugation and stored at -20°C for analysis. The BAL cells were washed with R10 medium (RPMI-1640 with 10% fetal bovine serum) before subsequent treatment or cryopreservation.

### ELISA assay to detect S1-specific antibody responses

The BAL samples were concentrated using Amicon Ultra centrifugal filter units (10kDa cutoff, *Millipore Sigma*), and the total IgG and IgA were determined using the Rhesus Monkey IgG-UNLB (*Southern Biotech*), and the Monkey IgA ELISA development kit (HRP) (MabTech) respectively, following the manufacturer’s protocol as described before ^20^. Nasal swab samples were put into 1 ml of 1XPBS buffer containing 0.1% BSA, 0.01% thimerosal, and 750 Kallikrein inhibitor units of aprotinin (Sigma) and stored at -80°C. Nasal swabs were thawed, and the recovered solution was passed through a 5 μm PVDF microcentrifugal filter unit (Millipore, Billerica, MA). The buffer flow-through was collected and stored at -20° until analysis.

ELISA assays were run as described before ^20^. The S1-specific binding assays were coated with 100 ng/well of the SARS-CoV-2 spike S1-His Recombinant Protein (Sino Biological) using high-binding 96-well plates (Santa Cruz Biotechnology). After incubation at 4°C overnight, and 1hr. blocking with 300 μL of 2% sodium casein in 1X PBS, the concentrated BAL samples (with a series of 2-fold dilutions starting from an IgA or IgG concentration of 2 μg/mL) or nasal swab samples, or serially diluted serum samples (4-fold starting from a 1:150 dilution) were applied in duplicate. After incubation at room temperature for 1 hr., the plates were washed four times. Subsequent steps of incubation with HRP-labeled secondary antibody and TMB substrate were followed as described before. For IgG and IgA binding assay, Goat Anti-Monkey IgG (alpha-chain specific)-HRP conjugate (1:5,000 dilutions, *Alpha Diagnostic*) and were used, respectively, as a secondary antibody. Area under the curve, endpoint titer, and ID50 values were calculated by GraphPad Prism 8 software with sigmoidal nonlinear regression. Dimeric IgA in BAL and nasal swabs was measured using DuoSet ELISA Ancillary Reagent Kit 2 (R&D Systems) as described before^20^. 100 ng/well of the SARS-CoV-2 spike S1 protein was coated and blocked. Original BAL samples or nasal swab flow-through from vaccinated and naïve animals were added in duplicate to the plates, followed by adding mouse anti-rhesus J chain [CA1L_33e1_A1a3] antibody (1:1000 dilutions, *NIH nonhuman primate reagent resource*), and Goat anti-mouse IgG-HRP conjugate (1:10,000 dilutions, *R&D Systems*). Each step was followed by 1 hr. incubation at room temperature and five washes.

### Plaque reduction neutralization test (PRNT)

The PRNT was performed in duplicate as described before ^20^. Vero E6 cells (ATCC, cat. no. CRL-1586), and 30 pfu challenge titers of SARS-CoV-2 virus USA-WA1/2020 strain or Vero TMPRSS2 cells (obtained from Dr. Adrian Creanga and Barney Graham, VRC, NIAID, Bethesda, MD) and same titer of the beta variant (B.1.351, SRA strain) was used to test the PRNT titers against the WA or beta variant of SARS-CoV-2 ^44^. Serum samples of 3-fold serial dilution starting from 1:20, and up to final dilution of 1: 4860 were incubated with 30 pfu of SARS-CoV-2 virus for 1 hr. at 37 °C. The serial dilutions of virus–serum mixtures were then added onto Vero E6 cell monolayers in cell culture medium with 1% agarose for 1 hr. at 37 °C with 5% CO_2_. The plates were fixed and stained after three days of culture. ID50 and ID90 were calculated as the highest serum dilution resulting in 50 and 90% reduction of plaques, respectively.

### Intracellular cytokine staining assay

SARS-CoV-2-specific T cells were measured from BAL and PBMC samples by flow cytometric intracellular cytokine analysis, as previously described ^20,45,46^. Briefly, 2 μg/ml of SARS-CoV-2 S1 protein (*Sino Biological*) for PBMC, and 5 μg/ml for BAL samples was incubated with cell samples at 37°C 5%CO_2_ overnight in the presence of 0.15 μg/ml of brefeldin A. Negative and positive controls were stimulated with medium-only (no S1 protein) or with cell activation cocktail with PMA (20.25 pM) and ionomycin (335 pM) and 0.15 μg/ml of brefeldin A (Biolegend). Cells were stained with viability dye (Invitrogen) and the following antibody mixtures: PE-Cy7-CD3, BV605-CD4, APC-Cy7-CD8, Alexa Fluor® 700-CD45 were from BD Biosciences, FITC-CD28, Pe-Cy5-CD95, BV711-TNFα, IFNγ-PE or - PerCP, Alexa Fluor® 647-IL4, BV785-IL2, BV421-IL-17A, BV785-CD14, BV421-CD16 were from Biolegend; PE-IL13 was from Miltenyi Biotech. Detailed antibody information is listed in the previous publication ^20^. Data acquisition and analyses were performed using an LSRII flow cytometer with 4 lasers (BD Bioscience) and FlowJo software (Becton Dickinson). The antigen-specific T cell responses were reported as the frequencies of cytokine-positive cells in the samples stimulated with S1 protein minus those in the medium-only control.

### SARS-CoV-2 beta variant viral challenge

Four weeks after the one-year boost, 5 vaccinated and 5 control animals were challenged with 1×10^5 pfu SARS-CoV-2 virus beta variant (seed stock obtained from BEI Resources; NR-54974, B.1.351, SRA strain). The challenge stock was grown in Calu-3 cells and was deep sequenced, which confirmed the expected sequence identity with no mutations in the Spike protein greater than >2.5% frequency and no mutations elsewhere in the virus at >13% frequency. The same beta variant stock was used in the earlier macaque challenge study at the same facility ^28^. To make sure that the virus was delivered to both upper and lower airway simultaneously, the diluted virus was given intranasally and intratracheally, each route with 1ml (0.5ml for each nostril). Nasal swab and BAL fluid samples were collected after challenge to measure the viral load.

### TCID50 assays to measure viral loads

Vero TMPRSS2 cells (obtained from the Vaccine Research Center-NIAID) were plated at 25,000 cells/well in DMEM + 10% FBS + Gentamicin and the cultures were incubated at 37°C, 5.0% CO2. Cells should be 80 -100% confluent the following day. Medium was aspirated and replaced with 180 μL of DMEM + 2% FBS + gentamicin. Twenty (20) μL of sample was added to top row in quadruplicate and mixed using a P200 pipettor 5 times. Using the pipettor, 20 μL was transferred to the next row, and repeated down the plate (columns A-H) representing 10-fold dilutions. The tips were disposed for each row and repeated until the last row. Positive (virus stock of known infectious titer in the assay) and negative (medium only) control wells were included in each assay set-up. The plates were incubated at 37°C, 5.0% CO2 for 4 days. The cell monolayers were visually inspected for CPE. Non-infected wells will have a clear confluent cell layer while infected cells will have cell rounding. The presence of CPE was marked on the lab form as a + and absence of CPE as -. The TCID50 value was calculated using the Read-Muench formula. For optimal assay performance, the TCID50 value of the positive control should test within 2-fold of the expected value.

### Histopathology and immunohistochemistry of lung sections

Seven days after SARS-CoV-2 viral challenge all the animals were necropsied and the lung tissue specimens were collected, fixed, processed, and embedded in paraffin blocks and sectioned at a thickness of 5 µm as described in the previous study ^20^. Briefly, hematoxylin and eosin (H&E) sections were examined under light microscopy and scored by a board-certified veterinary pathologist, who was blind to the groups. A rabbit polyclonal SARS-CoV-2 antibody (GeneTex) was used immunohistochemically to stain for the presence of SARS-CoV-2 virus antigen. An Olympus BX51 brightfield microscope was used, and representative photomicrographs were captured using an Olympus DP73 camera.

### Statistical analysis

Prism version 8 (Graph Pad) was used for statistical analyses. Area under curve (AUC) values were calculated for viral load, and Mann-Whitney tests were used for group comparisons as shown in the figures. A P value less than 0.05 was considered significant, and all statistical tests were 2-tailed.

### Study approval

Vaccination was performed at the National Institutes of Health NCI Animal Facility, Bethesda, MD, an American Association for the Accreditation of Laboratory Animal Care (AAALAC)-accredited facility with PHS Approved Animal Welfare Assurance (Assurance ID A4149-01). Animal Protocol No. VB-037 was approved by the NCI Animal Care and Use Committee (ACUC) to conduct the study. Two weeks before viral challenge, all 10 animals were moved to a qualified BSL3 facility at BIOQUAL, Inc.. The SARS-CoV-2 viral challenge study was approved and performed under BIOQUAL’s IACUC approved Protocol No. 20-107.

## Acknowledgments

This work was supported by funding from the Intramural Research Program, National Institutes of Health, National Cancer Institute, Center for Cancer Research funding Z01 BC-011941 to Jay A. Berzofsky. The NIH nonhuman primate reagent resource provided mouse anti-rhesus J chain antibody. The staff of the Laboratory Animal Sciences Program, Frederick National Laboratory for Cancer Research, provided technical support and animal care. The following reagent was deposited by the Centers for Disease Control and Prevention and obtained through BEI Resources, NIAID, NIH: SARS-Related Coronavirus 2, Isolate USA-WA1/2020, NR-53780 and Isolate beta variant B.1.351, NR-54974. We thank Shelby O’Connor for sequencing of the beta variant challenge stock.

The content of this publication does not necessarily reflect the views or policies of the Department of Health and Human Services, nor does mention of trade names, commercial products, or organizations imply endorsement by the U.S. Government.

## Author contributions

YS, JAB designed and interpreted the project. YS, JL processed samples, ran cellular assays. JL, TH, RZ, SP, YS performed antibody assays. LP performed PRNT assays. JT, YS prepared the PLGA nanoparticle, and other vaccines. IM, KB, MM, BMN performed pathology. HA, AC, RB, ET, JV, MB, JK led the animal studies. YS, JAB, HA, LL, ML, LW participated in study design and interpreted the experiments. DV, HC and YS performed statistical analyses. YS and JAB wrote the manuscript with input from all the coauthors.

## Notes

### Competing Interest Statement

The authors have declared no competing interest.

