## Supplementary material for "An intranasally administrated SARS-CoV-2 beta variant subunit booster vaccine prevents beta variant viral replication in rhesus macaques"

**Supplementary Figure 1.** T helper subsets in PBMC samples of the vaccinated macaques. The frequencies of IFNγ+, TFNα+, or IL4+ CD4$^{+}$ T cells were stained and measured after stimulation with PMA + ionomycin for 18hrs in PBMCs from different time-points.

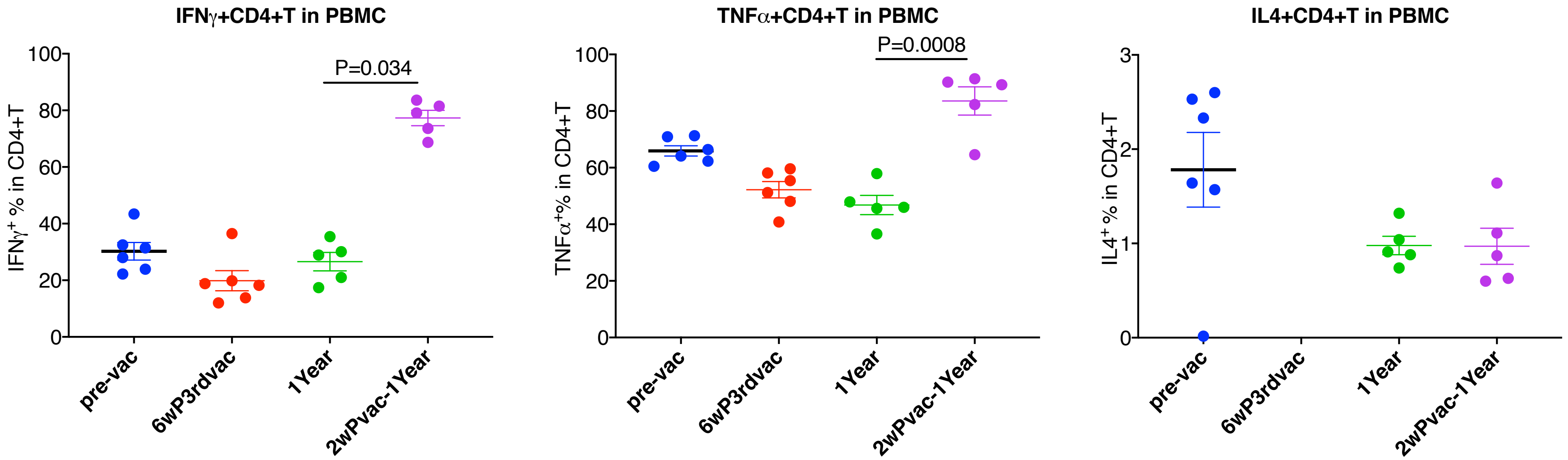

**Supplementary Figure 2.** Spearman's correlation among different types of humoral immune responses after the administration of one-year booster

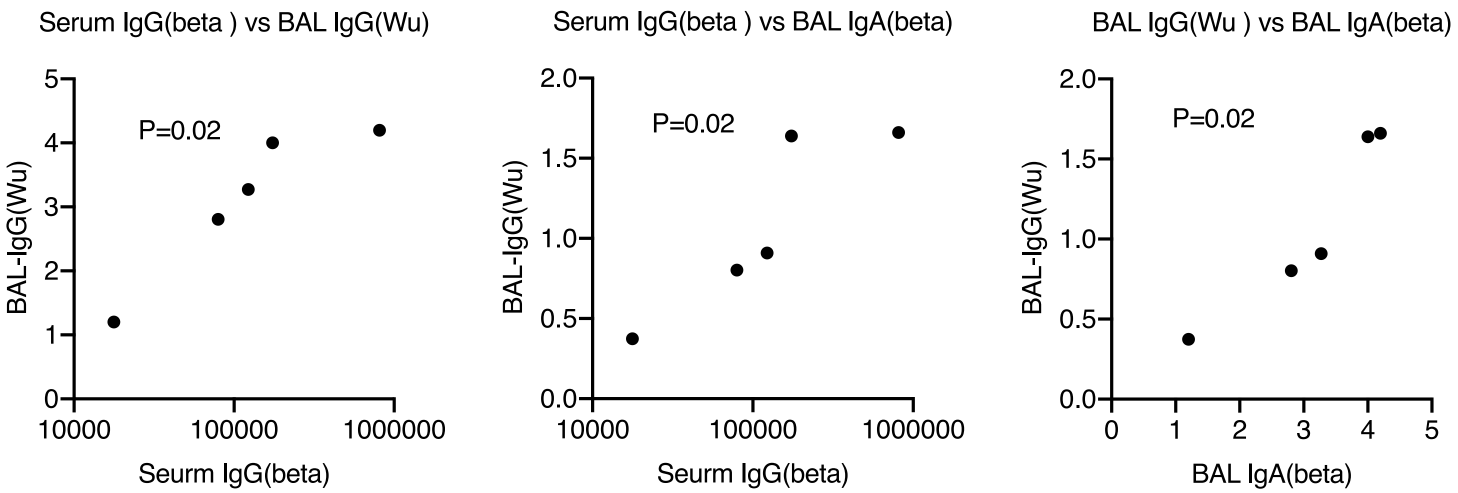

**Supplementary Table 1. Basic information of the animals enrolled in the study**

| Group | Animal ID | Sex | Date birth | Weight |
|---|---|---|---|---|
| Vaccine | DGVC | Male | 05/01/2016 | 5.7 |
| Vaccine | DGNi | Male | 03/18/2016 | 6.5 |
| Vaccine | DG1E | Male | 04/07/2016 | 6.8 |
| Vaccine | DGZT | Male | 04/04/2016 | 5.5 |
| Vaccine | DGLM | Male | 04/14/2015 | 5.1 |
| Control | 14D044 | Female | 06/19/2014 | 4.07 |
| Control | 14D064 | Female | 06/19/2014 | 4.8 |
| Control | 14D080 | Female | 08/08/2014 | 4.28 |
| Control | 14D065 | Female | 06/29/2014 | 4.68 |
| Control | 14D068 | Female | 07/15/2014 | 4.8 |

### Supplmentary Table 2. correlations among the humoral responses induced by one-year booster

| **Spearman P** | Serum IgG (Wu) | Seurm IgG(beta) | BAL-IgG(Wu) | BAL-IgG(beta) | PRNT50(WA) | PRNT50(beta) | PRNT90(WA) | PRNT90(beta) | Nasal IgA (Wu) | Nasal IgA (beta) | Nasal sIgA*(Wu) | Nasal sIgA(beta) | BAL IgA (Wu) | BAL IgA (beta) | BAL sIgA(Wu) |
|---|---|---|---|---|---|---|---|---|---|---|---|---|---|---|---|
| Serum IgG (Wu) | | | | | | | | | | | | | | | |
| Seurm IgG(beta) | 1.00 | | | | | | | | | | | | | | |
| BAL-IgG(Wu) | 1.00 | **0.02** | | | | | | | | | | | | | |
| BAL-IgG(beta) | 0.95 | **0.08** | **0.08** | | | | | | | | | | | | |
| PRNT50(WA) | 0.80 | **0.07** | **0.07** | **0.13** | | | | | | | | | | | |
| PRNT50(beta) | 1.00 | **0.10** | **0.10** | 0.30 | 0.20 | | | | | | | | | | |
| PRNT90(WA) | 0.67 | 0.47 | 0.47 | 0.80 | 0.37 | 0.20 | | | | | | | | | |
| PRNT90(beta) | 1.00 | **0.10** | **0.10** | 0.30 | 0.20 | **0.10** | 0.20 | | | | | | | | |
| Nasal IgA (Wu) | 0.68 | 0.23 | 0.23 | 0.35 | **0.13** | 0.30 | 0.33 | 0.30 | | | | | | | |
| Nasal IgA (beta) | 0.52 | 0.35 | 0.35 | 0.23 | 0.27 | 0.60 | 0.73 | 0.60 | **0.08** | | | | | | |
| Nasal sIgA(Wu) | 0.95 | **0.08** | **0.08** | 0.23 | **0.07** | **0.10** | 0.20 | **0.10** | **0.08** | 0.23 | | | | | |
| Nasal sIgA(beta) | 0.68 | 0.23 | 0.23 | 0.52 | 0.20 | **0.10** | **0.07** | **0.10** | **0.13** | 0.45 | **0.08** | | | | |
| BAL IgA (Wu) | 0.95 | **0.08** | **0.08** | **0.13** | 0.20 | **0.10** | 0.47 | **0.10** | 0.52 | 0.68 | 0.23 | 0.35 | | | |
| BAL IgA (beta) | 1.00 | **0.02** | **0.02** | **0.08** | **0.07** | **0.10** | 0.47 | **0.10** | 0.23 | 0.35 | **0.08** | 0.23 | **0.08** | | |
| BAL sIgA(Wu) | 0.68 | 0.78 | 0.78 | 0.95 | 0.53 | 0.60 | **0.13** | 0.60 | 0.23 | 0.52 | 0.35 | **0.13** | 1.00 | 0.78 | |
| BAL sIgA(beta) | 0.78 | 0.78 | 0.78 | 0.45 | 0.67 | 1.00 | 0.33 | 1.00 | 0.68 | 0.35 | 0.95 | 0.68 | 0.95 | 0.78 | 0.68 |

| **Spearman R** | Serum IgG (Wu) | Seurm IgG(beta) | BAL-IgG(Wu) | BAL-IgG(beta) | PRNT50(WA) | PRNT50(beta) | PRNT90(WA) | PRNT90(beta) | Nasal IgA (Wu) | Nasal IgA (beta) | Nasal sIgA(Wu) | Nasal sIgA(beta) | BAL IgA (Wu) | BAL IgA (beta) | BAL sIgA(Wu) |
|---|---|---|---|---|---|---|---|---|---|---|---|---|---|---|---|
| Serum IgG (Wu) | | | | | | | | | | | | | | | |
| Seurm IgG(beta) | 0.00 | | | | | | | | | | | | | | |
| BAL-IgG(Wu) | 0.00 | 1.00 | | | | | | | | | | | | | |
| BAL-IgG(beta) | 0.10 | 0.90 | 0.90 | | | | | | | | | | | | |
| PRNT50(WA) | -0.16 | 0.95 | 0.95 | 0.79 | | | | | | | | | | | |
| PRNT50(beta) | 0.22 | 0.89 | 0.89 | 0.67 | 0.83 | | | | | | | | | | |
| PRNT90(WA) | 0.26 | 0.53 | 0.53 | 0.16 | 0.56 | 0.83 | | | | | | | | | |
| PRNT90(beta) | 0.22 | 0.89 | 0.89 | 0.67 | 0.83 | 1.00 | 0.83 | | | | | | | | |
| Nasal IgA (Wu) | 0.30 | 0.70 | 0.70 | 0.60 | 0.79 | 0.67 | 0.58 | 0.67 | | | | | | | |
| Nasal IgA (beta) | 0.40 | 0.60 | 0.60 | 0.70 | 0.63 | 0.45 | 0.21 | 0.45 | 0.90 | | | | | | |
| Nasal sIgA(Wu) | 0.10 | 0.90 | 0.90 | 0.70 | 0.95 | 0.89 | 0.74 | 0.89 | 0.90 | 0.70 | | | | | |
| Nasal sIgA(beta) | 0.30 | 0.70 | 0.70 | 0.40 | 0.74 | 0.89 | 0.95 | 0.89 | 0.80 | 0.50 | 0.90 | | | | |
| BAL IgA (Wu) | 0.10 | 0.90 | 0.90 | 0.80 | 0.74 | 0.89 | 0.53 | 0.89 | 0.40 | 0.30 | 0.70 | 0.60 | | | |
| BAL IgA (beta) | 0.00 | 1.00 | 1.00 | 0.90 | 0.95 | 0.89 | 0.53 | 0.89 | 0.70 | 0.60 | 0.90 | 0.70 | 0.90 | | |
| BAL sIgA(Wu) | 0.30 | 0.20 | 0.20 | -0.10 | 0.37 | 0.45 | 0.79 | 0.45 | 0.70 | 0.40 | 0.60 | 0.80 | 0.00 | 0.20 | |
| BAL sIgA(beta) | 0.20 | -0.20 | -0.20 | -0.50 | -0.26 | 0.22 | 0.58 | 0.22 | -0.30 | -0.60 | -0.10 | 0.30 | 0.10 | -0.20 | 0.30 |

*sIgA: secretory IgA

**Supplementary Table 3. Inflammation and immunohistochemistry evaluation of the lung sections (necropsy at day 7 post SARS-CoV-2 challenge)**

| Animal ID | Group | H&E (Lc; Rm; Rc) | COVID-19 IHC (Lc; Rm; Rc) |
|---|---|---|---|
| DGVC | Vaccine | +/-; +/-; +/- | -; -; - |
| DGNi | Vaccine | +/-; +/-; + | -; -; - |
| DG1E | Vaccine | +/-; +/-; +/- | -; -; - |
| DGZT | Vaccine | +; + ; + | -; -; - |
| DGLM | Vaccine | +/-; +/-; +/- | -; -; - |
| 14D044 | Control | +/-; +; + | -; -; - |
| 14D064 | Control | ++; +-; ++ | -; -; + |
| 14D080 | Control | +/-; +; ++ | +; -; +/- |
| 14D065 | Control | +; + ; ++ | -; -; +/- |
| 14D068 | Control | +/-; +; + | -; +/-; - |

H&E (inflammation) severity scale: normal= - (0); <10% (tissue affected) = +/- (1); >10-<25% = + (2) ; >26-<50% = ++ (3); >50%= +++ (4). Three parts of the lung lobes: Left caudal [Lc], Right Middle [Rm], and Right caudal [Rc].

IHC (COVID + foci)
Scoring: - = no SARS-CoV-2 Ag detected; +/- = rare- occasional; + = occasional-multiple; ++ = multiple- numerous (foci often larger); +++ = numerous
